# A Transcriptional Cofactor Regulatory Network for the *C. elegans* Intestine

**DOI:** 10.1101/2023.01.05.522920

**Authors:** Brent B. Horowitz, Shivani Nanda, Albertha J.M. Walhout

## Abstract

Chromatin modifiers and transcriptional cofactors (collectively referred to as CFs) work with DNA-binding transcription factors (TFs) to regulate gene expression. In multicellular eukaryotes, distinct tissues each execute their own gene expression program for accurate differentiation and subsequent functionality. While the function of TFs in differential gene expression has been studied in detail in many systems, the contribution of CFs has remained less explored. Here we uncovered the contributions of CFs to gene regulation in the *Caenorhabditis elegans* intestine. We first annotated 366 CFs encoded by the *C. elegans* genome and assembled a library of 335 RNAi clones. Using this library, we analyzed the effects of individually depleting these CFs on the expression of 19 fluorescent transcriptional reporters in the intestine and identified 216 regulatory interactions. We found that different CFs interact specifically with different promoters, and that both essential and intestinally expressed CFs exhibit the highest proportion of interactions. We did not find all members of CF complexes acting on the same set of reporters but instead found diversity in the promoter targets of each complex component. Finally, we found that previously identified activation mechanisms for the *acdh-1* promoter use different CFs and TFs. Overall, we demonstrate that CFs function specifically rather than ubiquitously at intestinal promoters and provide an RNAi resource for reverse genetic screens.

## INTRODUCTION

Proper spatiotemporal gene expression is necessary for organismal growth and development, as well as maintaining cellular homeostasis and mounting stress responses. Gene expression depends on chromatin state and is governed on many levels. First and foremost, gene expression is regulated by transcription factors (TFs) that bind DNA elements located in gene promoters and enhancers and either activate or repress transcription. Most TFs regulate multiple genes, and each gene may be controlled by several TFs. Such complex relationships among genes and TFs can be captured in gene regulatory networks (GRNs)(Rogers and Bulyk 2018; Walhout 2006). Expression profiling by RNA-seq has enabled the identification of gene expression programs for many individual tissues and cells in a variety of organisms. However, the underlying GRNs that direct those programs are not well understood.

While TFs are often considered the primary drivers of specific gene expression programs, transcriptional cofactors (CFs) also play central roles in regulating transcription and therefore in establishing GRNs. CFs, which do not bind DNA directly, fall into two classes. The first class of CFs interact with TFs and RNA polymerase II at the promoter and regulate polymerase activity during several steps in the transcription cycle, including initiation, pausing, elongation and termination (Cramer 2019; Haberle and Stark 2018). The second class are chromatin modifiers that covalently modify histones or remodel nucleosomes to change chromatin structure and accessibility for TFs and RNA polymerase II (Bannister and Kouzarides 2011; Talbert et al. 2019).

CFs often function within multiprotein complexes that frequently show modularity. Smaller complexes tend to have core members all of which are essential for complex function and regulate the expression of similar gene sets, *i*.*e*., loss or perturbation of each disrupts complex functionality (Kemmeren et al. 2014). Larger complexes, such as the thirty-member Mediator complex, are often composed of submodules, each responsible for regulating the expression of subsets of genes (El Khattabi et al. 2019; Kemmeren et al. 2014; Lenstra et al. 2011). Additionally, submodules of some CF complexes, such as SAGA and Mediator, can work both together and independently of each other to regulate gene expression (Anandhakumar et al. 2016; Li et al. 2017). Further, subunits are sometimes shared between complexes. For example, the NuA4 histone acetyltransferase complex and the histone depositing complex SWR1-C share four components(Lu et al. 2009). This complexity makes it difficult to study how CFs influence the expression of individual genes, and how interactions with other CF complexes, TFs and the basal transcriptional machinery affect gene regulation.

Chromatin modifying enzymes and the epigenetic marks placed by some of these enzymes are found at specific positions throughout the genome, yet not every gene associated with these factors changes expression upon their depletion (Lenstra et al. 2011; Venters et al. 2011; Weiner et al. 2012). This indicates that CFs have specific effects on gene expression that cannot be solely explained by binding profiles. How then does this specificity arise and what conditions lead to CF-specific gene regulation? Recent work has shown that certain CF complexes that were once thought to universally regulate gene expression are instead specific for certain types of promoters and enhancers (Haberle et al. 2019; Neumayr et al. 2022). Additionally, some chromatin remodeler complexes specify spatiotemporal gene expression throughout development, while others are used to regulate housekeeping genes that have stable developmental expression (Hendy et al. 2022). This indicates that nucleosome/chromatin structure may be different at these regulatory regions requiring different types of remodeler activities for gene activation and/or repression.

Delineating how CFs work together and with TFs to establish gene specificity in various contexts is important for understanding GRNs and how these influence growth, development, homeostasis, and cellular reactions to changing environments. Strategies for delineating GRNs have fallen into two categories: regulator (or protein)-centered approaches and gene-centered approaches (Araya et al. 2014; Fuxman Bass et al. 2016; Fuxman Bass et al. 2015; Johnson et al. 2007; Kemmeren et al. 2014; MacNeil et al. 2015; McIsaac et al. 2012). Regulator-centered approaches identify the global complement of genes bound and/or regulated by individual factors while gene-centered approaches focus on a specific gene and its regulatory sequences to identify the complement of regulators that bind or regulate the activity of that gene.

Determining GRNs for specific tissues or cells has been challenging. Single-cell RNA-seq has enabled gene expression measurement at the level of individual cells, and these measurements can be grouped into different cell and tissue types (Dixit et al. 2016). Other studies have inferred tissue-specific GRNs from a combination of gene expression data and protein-protein interactions (Sonawane et al. 2017). However, it is only just becoming feasible to systematically perturb each TF and measure tissue-specific gene expression changes at the same resolution in order to elucidate function (Adamson et al. 2016; Dixit et al. 2016; Liu et al. 2022).

The nematode *Caenorhabditis elegans* provides a powerful model to delineate GRNs for individual tissues. *C. elegans* has a transparent body, which allows the visualization of fluorescent proteins expressed under the control of specific gene promoters *in vivo*(Chalfie et al. 1994). With *C. elegans* transgenes, changes in fluorescent reporter expression in specific tissues upon gene knockdown can be monitored visually in living animals. This approach was used with a collection of 19 transgenic strains and comprehensive TF RNAi to delineate an *in vivo* activity-based GRN for the *C. elegans* intestine. This GRN contains 411 regulatory interactions driven by 177 TFs (MacNeil et al. 2015). TF knockdown mostly decreased reporter expression, indicating that TFs overall predominantly function as activators. One major insight from this study was that many of the interactions detected are likely indirect. Many TFs that showed activity in the screen were not found to physically interact with the promoters they regulate. Organizing the effects of different TFs on different promoters by nested effects modeling led to a hierarchical model in which TFs, directly or indirectly, regulate other TFs. Importantly, the TFs that were placed low in the hierarchy tended to physically interact with the promoters they regulate.

The *C. elegans* intestine is a highly metabolic tissue that functions not only as gut, but also as liver (Kaletsky et al. 2018; Yilmaz et al. 2020). Since gene expression is frequently regulated by changes in metabolism and *vice versa* (Giese et al. 2019; Watson et al. 2015), a similar RNAi screen to that above was performed by depleting ∼1,500 metabolic genes and testing their effect on the activity of the same 19 promoters. This led to the identification of 1,251 regulatory interactions involving 512 metabolic genes (Bhattacharya et al. 2022). In contrast to TF knockdown, metabolic gene knockdown tended to increase reporter expression, indicating that metabolic perturbations mostly activate transcription. Interestingly, it was found that certain types of metabolism affect promoter activity over other types. For instance, many promoters were affected by perturbations in oxidative phosphorylation, while little to no effect was seen upon depletion of carbohydrate metabolism. Additional insights were gained through the identification of TFs that act downstream of some of these metabolic processes. Overall, the above results examining the interplay between TFs and metabolic enzymes in the regulation of gene expression suggest that in the *C. elegans* intestine, metabolic perturbations are commonly sensed by TFs that then function either directly or indirectly to regulate gene expression.

Since TFs work with non-DNA binding CFs to activate or repress their targets, we reasoned that additional information about the GRN in the *C. elegans* intestine could be obtained by asking globally how CFs influence promoter activity. While previous studies have individually deleted the entire complement of yeast CFs or degraded specific mammalian CFs and examined transcriptome-wide changes in gene expression (Haberle et al. 2019; Hendy et al. 2022; Lenstra et al. 2011; Neumayr et al. 2022), no study has comprehensively depleted CFs and examined the effects on a per gene basis. Here, we extend our gene-centered approach of using RNAi and specific promoters driving fluorescent reporters to screen a library of transcriptional CFs. We assembled a library of RNAi strains targeting most CFs encoded by the *C. elegans* genome and examined the effects of those CF knockdowns on the same 19 strains that were assessed in previous studies (Bhattacharya et al. 2022; MacNeil et al. 2015). We found that, while most CFs had no effect on the 19 promoter reporters, a few CFs affected many promoters. Additionally, we assessed which CFs function in three independent activation mechanisms of a single promoter reporter, *Pacdh-1::GFP*. This study provides a basis for better understanding how CFs function within the *C. elegans* intestine as well as an RNAi resource for future studies.

## MATERIALS AND METHODS

### CF annotations

Previously, a preliminary set of CF predictions used the following categories: histone methyltransferases, histone demethylases, histone acetyltransferases (HATs), histone deacetylases (HDACs), TATA-binding protein associated factors, Mediator components, and any gene encoding a protein with a plant homeodomain, chromodomain, or bromodomain(Reece-Hoyes et al. 2013). There was also a set of literature-defined CFs in an ‘other’ category. We eliminated genes previously annotated as CFs that had no annotated CF function but only encoded enzymes with a particular enzymatic function (for example, a cytosolic methyltransferase with no histone targets)(Reece-Hoyes et al. 2005; Reece-Hoyes et al. 2013). We also removed reannotated pseudogenes and dead genes. We then added categories for chromatin remodelers, histone kinases, histone phosphatases, histone ubiquitinases, coregulators, RNA polymerase II-associated factors, and Tudor-domain containing proteins. Using WormBase version WS284, we searched for additional genes matching those categories Gene Ontology and InterPro protein motif terms (Ashburner et al. 2000; Blum et al. 2021). Additional factors were found through homology to yeast and human CFs. We also revised the literature-defined CFs in the other category, including adding newly annotated CFs acting in dosage compensation (Brejc et al. 2017). In total, we annotated 366 *C. elegans* CFs (**Table S1)**.

### Essentiality enrichment analysis

*C. elegans* phenotypes were obtained using the SimpleMine tool in WormBase version WS284 (http://wormbase.org). A total of 19,987 protein-coding genes were included in the analysis. We considered essential genes as any gene with at least one of the following phenotypes: lethal, larval lethal, larval arrest, embryonic lethal, embryonic arrest, or sterile. Hypergeometric distribution was used to determine which CF categories or complexes are enriched for essential phenotypes **(Table S2)**.

### Intestinal expression of CFs

We previously used a single-cell RNA-seq dataset to derive tissue expression of *C. elegans* genes at the second larval stage (L2)(Yilmaz et al. 2020). Here, we used this tissue gene expression data to evaluate intestinal expression of CFs.

### CF RNAi library

To construct the CF RNAi library, we obtained all available RNAi clones in from either the ORFeome or the Ahringer library (Kamath et al. 2003; Rual et al. 2004). We verified each clone by Sanger sequencing and if both clones for a given gene were correct, we only included the ORFeome clone. If a CF was not available in either library, we attempted to clone the ORF from newly synthesized cDNA (see below). The Gateway system was used to clone each cDNA into pDONR221, then subsequently into L4440-Dest-RNAi vector, followed by transformation into the RNAi-competent *E. coli* strain HT115 (Walhout et al. 2000). All new clones were sequence verified. The final library was organized first by function, then alphabetically in 96-well plates; each plate also included vector control, GFP and mCherry RNAi clones, and blank wells (**Table S1**). The library contains 335 RNAi clones: 186 from the ORFeome, 95 from the Ahringer library, and 54 new clones. RNAi clones for *tag-153* and *cbp-2* are not included in the library plates and were cloned separately. Genotyping primers and primers for the *de novo* clones are listed in **Table S6**.

### *C. elegans* strains

*C. elegans* strains were maintained on nematode growth media (NGM) agar seeded with an *E. coli* OP50 diet as described, with some exceptions as noted below (Brenner 1974). The 19 transgenic promoter strains were described previously (**Table S3**)(MacNeil et al. 2015). The *mthf-1(ww50); nhr-10(tm4695); Pacdh-1::GFP* strain (Giese et al. 2020) was maintained on NGM using soy peptone in place of bactopeptone containing 0.64 nM vitamin B12 (Sigma V2876).

### RNAi Screening

RNAi screening was performed as previously described (Conte et al. 2015; MacNeil et al. 2015). Briefly, *E. coli* HT115 harboring individual CF RNAi plasmids were incubated overnight in 1 mL lysogeny broth (LB) + 50 μg/mL ampicillin + 10 μg/mL tetracycline in 96-deep-well plates at 37°C shaking at 200 rpm. The next day, 50 μL of overnight culture was transferred into 1 mL LB + 50 μg/mL ampicillin in 96-deep-well plates and incubated at 37°C at 200 rpm for six hours. Ten μL of this culture was added to 96-well plates of NGM containing 50 μg/mL ampicillin and 2 mM isopropyl ß-D-thiogalactopyranoside (IPTG). Plates were dried and incubated overnight at room temperature. 20 to 40 synchronized L1 animals were added to the bacteria containing NGM agar plates and incubated at 20°C for 48 hours. Animals exposed to CF RNAi were visually scored for changes in intestinal fluorescence relative to vector control animals. Fluorescence changes in other tissues were not recorded. Each of the 19 strains was screened three times verses the entire RNAi library. Primary hits were considered as any interaction which occurred in two or three of the three replicates.

Primary hits were retested as above in duplicate, using 24-well NGM agar plates with 50 μL of bacterial culture in each well and 80-100 animals per well. After this step hits were defined as any CF clone that scored positively in three of five combined replicates. Finally, RNAi was performed as above, using 6 cm plates, 100 μL of bacterial culture, and 150-200 animals per well. After 48h of animal growth, animals were photographed under brightfield and either GFP or mCherry channels. Final hits were those that were confirmed in at least one of two photographed replicates. The *tag-153* RNAi strain was tested alongside the remainder of the library and is included in the interaction analyses. The *cbp-2* RNAi strain was only tested in the final experiment and is not included in interaction counts but is included in photos in the supplementary tables.

### Promoter interaction enrichments

Hypergeometric distribution was used to determine CF categories or complexes that are enriched for either increases or decreases in fluorescence with each of the 19 transgenic promoter strains **(Table S5)**.

### qRT-PCR

*C. elegans* animals were grown as described above for RNAi screening using 10 cm NGM plates. After 48 hours of growth, animals were washed in M9 buffer and mRNA was extracted using the Direct-Zol RNA Miniprep Kit (Zymo Research R2050), including DNAse I treatment. cDNA was prepared using Oligo(dT) 12-18 Primer (Thermo Fisher 18418) and M-MuLV Reverse Transcriptase (NEB M0253). qPCR was performed in biological and technical triplicate using an Applied Biosystems QuantStudio 3 Real-Time PCR System with Fast SYBR Green Master Mix (Thermo Fisher 4385617). Relative transcript abundance was calculated using the ΔΔCt method (Livak and Schmittgen 2001). Endogenous controls were *act-1* and *ama-1*. Primer sequences are provided in **Table S6**.

### *Pacdh-1* activation mechanisms

For all three *Pacdh-1::GFP* activation conditions, we grew vector control, GFP RNAi, and CF or TF RNAi strains as above and plated as above. To test which TFs and CFs contribute to the regulation of the *acdh-1* promoter by vitamin B12 mechanism I (Bulcha et al. 2019), we incubated *Pacdh-1*::*GFP* animals on NGM agar containing 50 μg/mL ampicillin, 2 mM IPTG, 40 mM propionate, and 5 nM vitamin B12. To test which TFs and CFs contribute to the regulation of the *acdh-1* promoter by vitamin B12 mechanism II (Giese et al. 2020), we incubated *(wwls24[Pacdh-1::GFP, unc-119(+)]; nhr-10(tm4695); mthf-1(ww50)* animals on soy peptone-based NGM agar containing 50 μg/mL ampicillin, 2 mM IPTG, and 5 nM vitamin B12. To test which TFs and CFs contribute to the regulation of the *acdh-1* promoter by succinate dehydrogenase inhibition, referred to here as mechanism III, we incubated *sdha-2(tm1420); wwls24[Pacdh-1::GFP, unc-119(+)]* animals on NGM agar containing 50 μg/mL ampicillin, 2 mM IPTG, and 6.4 nM vitamin B12. For all three conditions, we photographed animals at 48h post plating as above.

### Microscopy

All fluorescence microscopy was performed on a Nikon Eclipse 90i with a Nikon DS-FI1 color camera, using NIS Elements software. Animals were washed off the plates with M9 buffer and paralyzed in 1mM levamisole in a microfuge tube. The suspended animals were briefly centrifuged, and 2 μL of paralyzed animals were placed on an agar pad. Animals were aligned with a hair pick and photographed with a 10X Nikon CFI Plan Fluor. GFP excitement range was 450-490 nm and emission was 500-550 nm, while mCherry excitation was from 528-553 nm and emission was 590-650 nm. Pseudocolorization was added linearly by NIS Elements, and images were not altered any further.

## RESULTS

### *C. elegans* CF annotation

We previously annotated 228 *C. elegans* CFs (Reece-Hoyes et al. 2013). To extend this analysis and to comprehensively predict the complement of CFs encoded by the *C. elegans* genome, we used a combination of Gene Ontology, InterPro motifs, and sequence homology to known CFs in other species (Ashburner et al. 2000; Blum et al. 2021). We classified CFs using the following categories: histone modifiers (methyltransferases, demethylases, histone acetyltransferases (HATs), histone deacetylases (HDACs), histone ubiquitinases, kinases, and phosphatases), chromatin remodelers, RNA polymerase II-associated factors, Mediator components, coactivators and corepressors, TATA binding protein-associated factors, DNA methylation enzymes, as well as proteins with plant homeodomains, chromodomains, bromodomains, or Tudor domains. We also included sixteen proteins that were identified as CFs without a specific molecular function or activity or defined as CFs in the literature and included these in an ‘other’ category. Overall, we annotated 366 *C. elegans* CFs (**Figure 1A, Table S1**).

**Figure 1.**
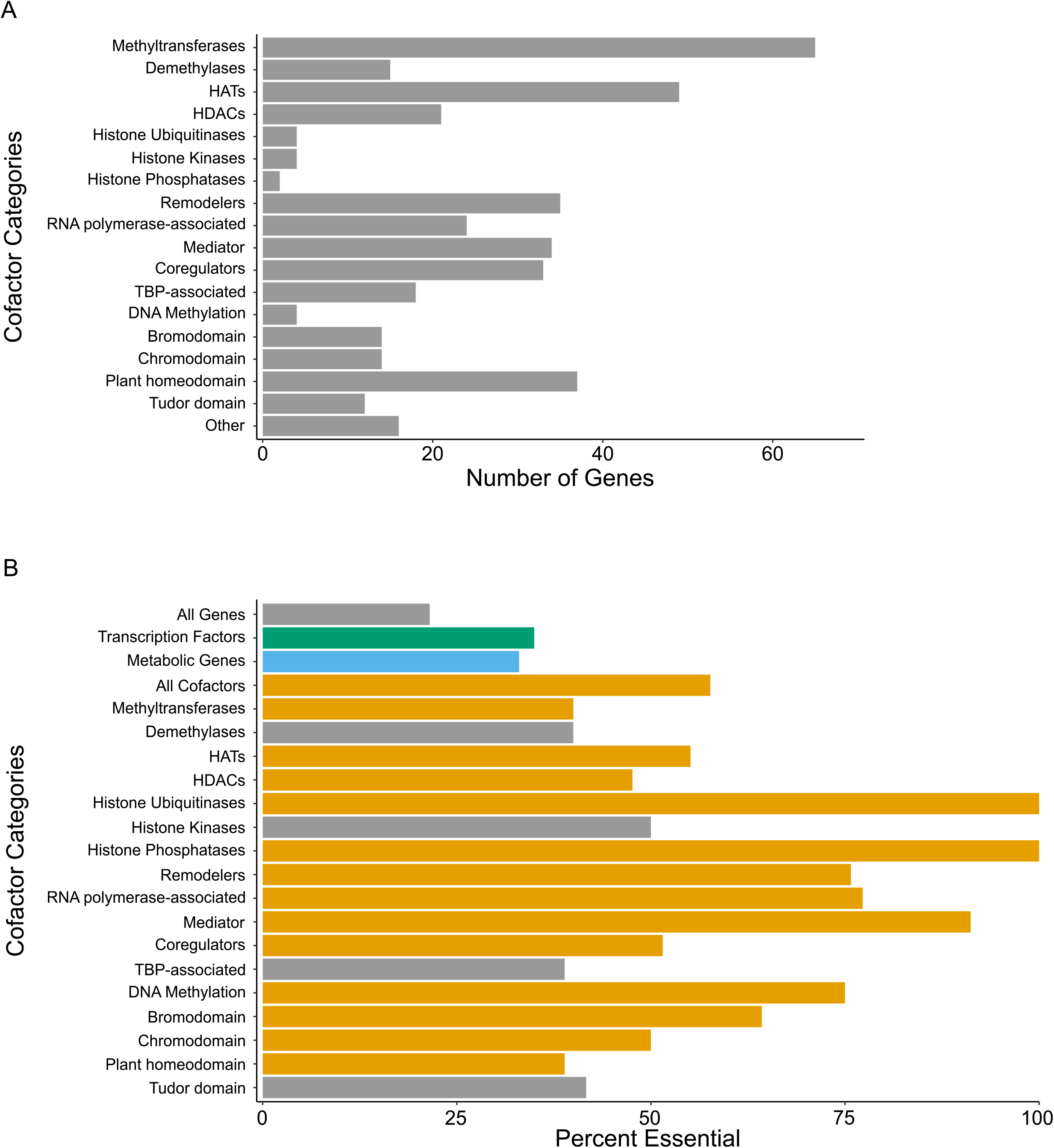
A compendium of *C. elegans* CFs. (A) Classification of the 366 CFs encoded by the *C. elegans* genome. Some CFs are counted in multiple categories; for instance, SET-8 is counted twice because it is a methyltransferase and has a plant homeodomain. (B) Percentage of each CF category that is annotated as essential (see Methods for essentiality definitions; P-values are listed in **Table S2**). Grey bars indicate no enrichment for essentiality. Green and blue bars indicate an essentiality enrichment for TFs and metabolic genes, respectively. Orange bars indicate CF categories that are enriched for essentiality.

Many CFs function in multiprotein complexes. We used Gene Ontology annotations to identify ten *C. elegans* CF complexes based on sequence homology with well-studied CFs in other organisms. These CF complexes are predicted to be comprised of three to 31 proteins (**Table 1**). When we compared the composition of the complexes to their human homologs, we found that we were able to identify all or nearly all *C. elegans* orthologs for each complex. The one exception was the SAGA complex where we were only able to identify half of the components (9/18). Some CFs are known or predicted to be present in multiple CF complexes. For example, EKL-4 is a component of both NuA4 and SWR1 complexes (Lu et al. 2009). For the Mediator complex, we used existing annotations of homology, as we could not use direct sequence comparisons to identify orthologs. Many Mediator complex components have limited sequence homology among species but do share certain common motifs (Bourbon 2008). The Mediator complex has low sequence conservation yet high structural conservation across metazoans, thus sequence comparisons alone are insufficient to identify those components (Bourbon 2008; Cai et al. 2009).

**Table 1.** *C. elegans* CF complexes. Names of the CF complexes are in the first column. Gene names are in the second column. * indicates not included in the CF RNAi library. Human orthologs are in the third column in the same order as the *C. elegans* gens in the second column. The fourth column contains any other complex members in humans we did not find in *C. elegans*.

| CF Complex | *C. elegans* Complex Components | Human Ortholog | Human Orthologs Not Found |
| --- | --- | --- | --- |
| CAF-1 | *chaf-1, chaf-2, rba-1* | *CHAF1A, CHAF1B, CHAF1C* |  |
| CCR4-NOT | *ccf-1, ccr-4, let-711, ntl-2, ntl-3, ntl-4, ntl-9, ntl-11, tag-153* | *CAF1, CCR4, CNOT1, CNOT2, CNOT3, CNOT4, CNOT9, CNOT11, CNOT2* | *CNOT10* |
| DRM | *dpl-1\*, efl-1\*, lin-9\*, lin-35, lin-37\*, lin-52, lin-53,* | *DP1, E2F2, LIN9, RBL2, LIN37, LIN52,* |  |
|  | *lin-54\** | *RBBP7, LIN-54* |  |
| Mediator | *cdk-4, cdk-8, cic-1\*, dpy-22, dyf-18, let-19, let-49, lin-25\*, mdt-4, mdt-6\*, mdt-8, mdt-10, mdt-11, mdt-15, mdt-17, mdt-18, mdt-19, mdt-20, mdt-21, mdt-22, mdt-27, mdt-29, mdt-31, R09F10.3\*, rgr-1, sna-1, sop-3, sur-2* | *CDK4, CDK19, CCNC, MED12L, CDK7, MED13L, MED7, MED24, MED4, MED6, MED8, MED10, MED15, MED17, MED18, MED19, MED20, MED21, MED22, MED27, MED29, MED31, MED27, MED14, MED6, MED1, MED23* | *MED16, MED25* |
| NuA4 | *B0025.4, cec-7, ekl-4, epc-1, F59E12.1, gfl-1, ing-3, mrg-1, mys-1, ruvb-1, ruvb-2, swsn-6\*, trr-1, Y43H11AL.1, ZK1127.3* | *MEAF6, MORF4L1, DMAP1, EPC1, BRD8, YEATS4, ING3, MORF4L1, KAT5, RUVBL1, RUVBL2, ACTL6B, TRRAP, ING3, MRGBP* | *MEAF6* |
| NuRD | *dcp-66, egl-27, hda-1, let-418, lin-40* | *GATAD2B, RERE, HDAC2, CHD3, MTA3* | *MBD2* |
| SAGA | *ada-2, pcaf-1, T22B7.4, taf-6.1, taf-6.2\*, taf-9, taf-10, taf-12, trr-1, Y17G9B.8\** | *TADA2B, KAT2B, SGF29, TAF6, TAF6, TAF9B, TAF10, TAF12, TRRAP, SGF29* | *ATXN7, ATXN7L3, ENY2, SF3B3, SF3B5, SUPT7L, SUPT20H, TADA1, TADA3, USP22* |
| Set1C/COMPASS | *ash-2, cfp-1, dpy-30, hcf-1, rbbp-5, set-2, swd-2.1, swd-2.2, wdr-5.1* | *ASH2L, CXXC1, DPY30, HCFC1, RBBP5, SETD1A, WDR82, WDR82, WDR5* |  |
| SWI/SNF/BAF | *C52B9.8, dpff-1, ham-3, let-526, snfc-5, swsn-1, swsn-2.2, swsn-3, swsn-4, swsn-6\*, swsn-7, ZK973.9* | *DPF2, SMARCD3, ARID1A, SMARCB1, SMARCC2, SMARCD1, SMARCE1, SMARCA2, ACTL6B, ARID2, SS18* |  |
| SWR1 | *arp-6\*, C17E4.6, ekl-4, F13C5.2\*, ruvb-1, ruvb-2, ssl-1, zhit-1* | *ACTR6, VPS72, DMAP1, BRD8, RUVBL1, RUVBL2, EP400, ZNHIT1* |  |

### CFs are enriched for essentiality

Previous work has demonstrated that *C. elegans* CFs often genetically interact with many other genes and pathways (Lehner et al. 2006). We therefore next asked whether *C. elegans* CFs tend to be essential for viability. We first used publicly available data mined from WormBase version WS284 to comprehensively define essential *C. elegans* genes (Lee et al. 2018). We then compared the percentage of genes that are essential among all *C. elegans* genes with the percentage of CF, TF, and metabolic genes, and found all three of these gene categories and many CF categories are significantly enriched for essentiality (**Figure 1B, Supplemental Table S2**). While methyltransferases, HATs, HDACs, histone ubiquitinases, histone phosphatases, remodelers, RNA-polymerase II-associated factors, Mediator components, DNA methylation enzymes and bromodomain-containing, chromodomain-containing, and plant homeodomain-containing proteins are more essential than random gene sets; demethylases, histone kinases, TATA binding protein-associated factors, and proteins harboring Tudor domains are not.

### A CF RNAi library resource

RNAi-by-feeding is a useful tool to examine phenotypes caused by gene knockdown in *C. elegans*. Previous work has established genome-wide RNAi libraries as well as libraries specific to TFs and metabolic genes (Bhattacharya et al. 2022; Fraser et al. 2000; Kamath et al. 2003; MacNeil et al. 2015; Rual et al. 2004). While the genome-wide libraries contained RNAi clones for many CFs, the strains are distributed among dozens of plates and many CFs are not represented in the libraries. In order to assess the effects of CF knockdowns, we assembled a library of RNAi strains targeting 335 of the 366 *C. elegans* annotated CFs (92%) (**Figure 2A, Table S1**). Of these, 186 clones were obtained from the ORFeome collection(Rual et al. 2004), 95 were retrieved from the Ahringer collection (Kamath et al. 2003), and 54 were generated *de novo*. All clones were sequence-verified. We arrayed all clones in four 96-well plates by function (**Figure 1A, Table S1**). Each plate contains vector controls as well as GFP and mCherry RNAi clones. This library is, to the best of our knowledge, the most comprehensive *C. elegans* CF RNAi resource.

**Figure 2.**
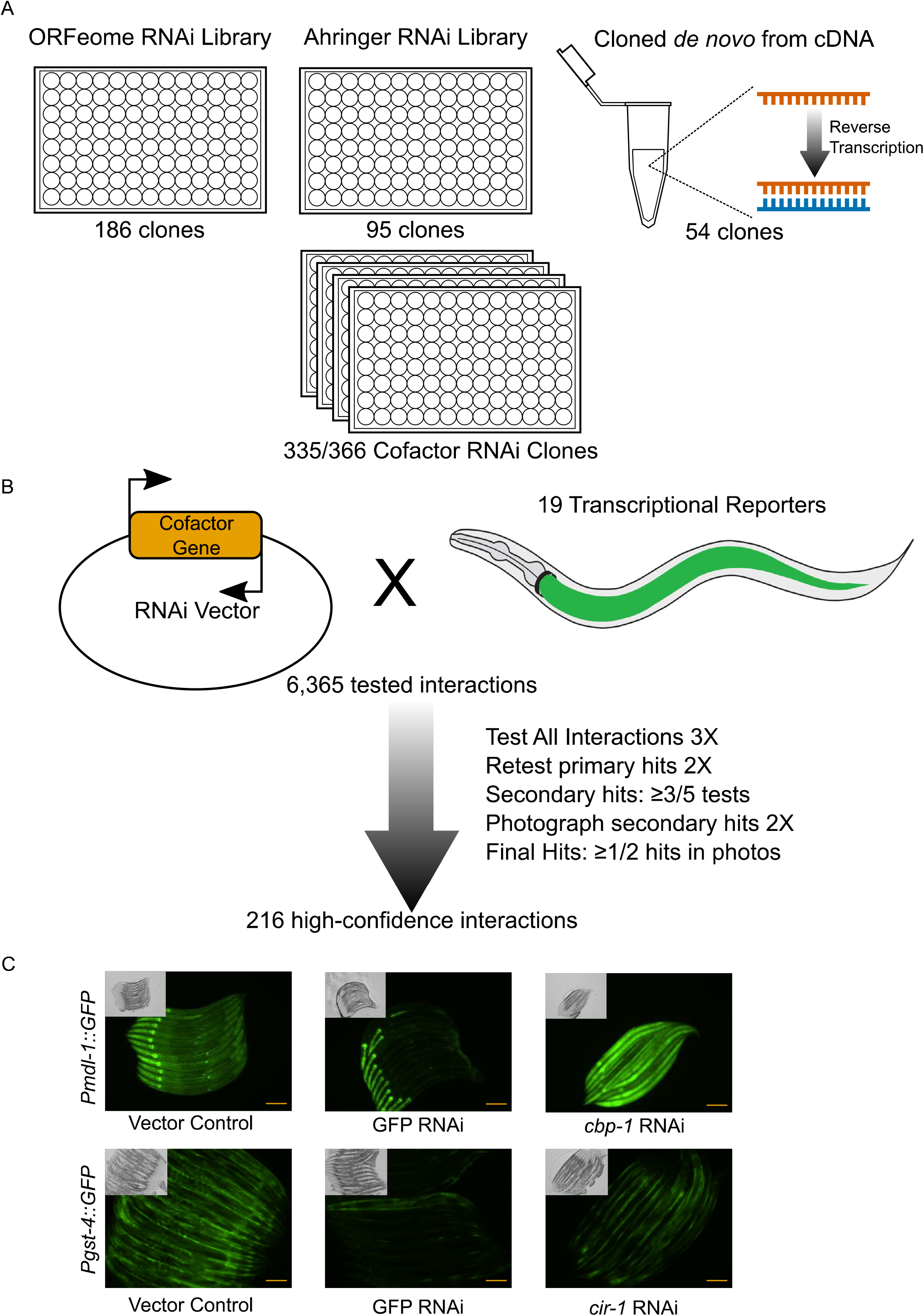
CF RNAi library assembly and RNAi screens. (A) Assembly of the CF RNAi library from the ORFeome and Ahringer RNAi libraries as well as clones generated *de novo*. (B) Diagram of the RNAi screening strategy. The primary screen was performed in triplicate and all hits were retested in duplicate. Any hits found in 3/5 tests were photographed twice; final hits were those that retested in at least one photograph. (C) Examples of CF RNAi changing intestinal fluorescence with two of the 19 promoter reporters. Scale bar=100μM.

### Uncovering regulatory promoter-CF interactions in the *C. elegans* intestine

To understand the regulatory effects of CFs on promoter activity, we performed a visual screen using 19 transcriptional reporter strains that express a fluorescent protein within the *C. elegans* intestine (MacNeil et al. 2015)(**Table S3**). We knocked-down expression of each CF by RNAi in each of the 19 reporter strains and visually monitored fluorescence changes in the intestine. This primary screen was done three times. Visual screens are noisy and many of the regulatory interactions found in the primary screen were subtle. Therefore, we retested any CF RNAi that caused a change in intestinal fluorescence in two or more replicates in any one reporter strain twice more with all 19 reporter strains using larger plates and more animals. Finally, we captured images for any interaction that was observed in at least three of the five tests of a given strain. Our final dataset includes only those interactions confirmed by a captured image (**Figure 2B-C, Figures S1-19 in File S1, Table S4**). Note that hits that are subtle yet consistent, such as *ntl-2* RNAi, which decreased intestinal fluorescence in the *Psbp-1::GFP* strain, were kept in the final dataset (**Figure S16 in File S1**).

Altogether, we detected 216 regulatory interactions between 19 promoters and 89 CFs (3.3% of all tested interactions) (**Figure 3A**). RNAi of more than half (61%) of these CFs showed a decrease in the expression of the intestinal reporter gene and therefore, these CFs potentially act as activators. Knockdown of the CBP/p300 HAT *cbp-1* changed the expression driven by 14 of the 19 promoters; ten reporters increased, and four reporters decreased in fluorescence. Therefore, *cbp-1* may function in both activation and repression of gene expression depending on the context. Alternatively, some of the observed interactions may be indirect. Depletion of most of the other CFs affected expression from only a small subset of the 19 promoters, indicating that these CFs mostly act in a gene-specific manner. We also observed an enrichment for interactions among essential CFs over non-essential CFs (**Figure 3B**).

**Figure 3.**
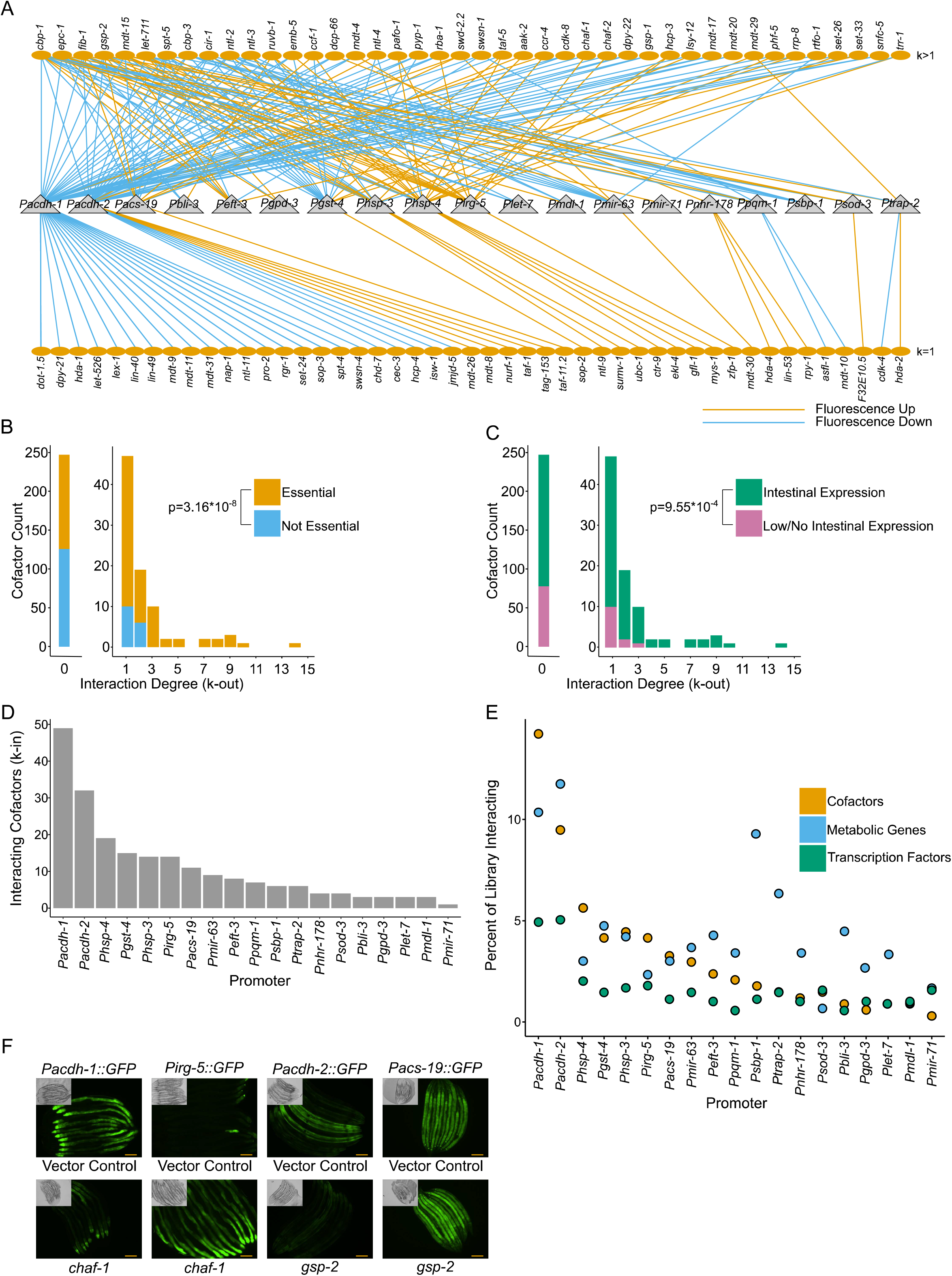
A CF regulatory network for the *C. elegans* intestine. (A) Bipartite graph of CF-promoter interactions. Triangles represent the 19 promoters. Ovals represent CFs regulating these promoters in the *C. elegans* intestine. Orange edges represent increases in fluorescence (repressing interactions) and blue edges represent decreases in fluorescence (activating interactions). k indicates the out-degree, or number of interactions, of each CF. (B) The k-out distribution. Orange bars represent essential CFs, blue bars represent CFs not annotated as essential. P-value was calculated using Mann-Whitney test. (C) As in B, green represents CFs expressed in the larval intestine, pink represents CFs not expressed in the larval intestine. P-value was calculated using Mann-Whitney test. (D) In-degree (k-in) for each of the 19 promoter reporters, or the number of CFs affecting each promoter. (E) Percentage of CFs, metabolic genes, and TFs (orange, blue, and green respectively) that interact with each promoter reporter, arranged by percent interacting CFs. (F) Examples of CFs that both increase and decrease fluorescence in different promoter reporters. Scale bar=100μM.

We next asked whether the CFs that affect any of the 19 promoters are expressed in the intestine or whether they are expressed in other tissues and therefore may exert their regulatory effects indirectly in a cell nonautonomous manner. We examined CF expression in a single-cell RNA-seq dataset that measured gene expression in seven tissues in the second larval (L2) stage and found that most *C. elegans* CFs are expressed in the animal’s intestine (**Figure 3C**)(Cao et al. 2017). We found an enrichment of intestinally expressed CFs for regulatory interactions, which indicates that the majority of the interactions uncovered likely act cell autonomously (**Figure 3C**).

To integrate our data with the TF GRN and metabolic MRN obtained in our previous screens we then looked for commonalities between TFs, metabolic genes, and CFs in their interactions with each promoter (Bhattacharya et al. 2022; MacNeil et al. 2015). Overall, we did not observe a uniform pattern. However, we did notice that the *Pacdh-1* and *Pacdh-2* promoters interacted with more CFs, TFs, and metabolic genes than other promoters (**Figure 3D-E, Figures S1-2 in File S1**). Other promoters, such as *Psbp-1* and *Ptrap-2*, were affected by depletion of many metabolic genes, but relatively few TFs or CFs. By contrast, the *hsp-4* promoter was affected by depletion of a relatively large proportion of CFs but few TFs and metabolic genes (**Figure 3E**).

Many CFs have been shown to have both activator and repressor functions in other organisms (Hunt et al. 2022; Subramaniam et al. 1999; Villanueva et al. 2011). Our data suggest that this may also be the case for *C. elegans*. In addition to *cbp-1* as discussed above, *chaf-1* RNAi decreased intestinal fluorescence of the *Pacdh-1::GFP* reporter, but increased GFP expression of the *Pirg-5::GFP* reporter (**Figure 3F**). Similarly, *gsp-2* RNAi had opposite effects on *Pacdh-2::GFP* and *Pacs-19::GFP* (**Figure 3F**). Overall, we found that, of the 42 CFs that interacted with at least two promoters, 27 both increased and decreased reporter expression when depleted (64%). This is modestly higher than the 50% of TFs and 41% of metabolic genes with multiple interactions that both activate and repress promoter activity (Bhattacharya et al. 2022; MacNeil et al. 2015). CF knockdowns also caused a higher proportion of fluorescence increases than TF knockdowns, but far less than metabolic gene knockdowns.

### CF complexes exhibit specificity and modularity

Since CFs function in large complexes and many have overlapping functions, we asked whether CF complex components or CFs in the different functional categories interacted with the same set of promoters. Four CF categories and members of eight different complexes were enriched for interactions with at least one promoter (**Figure 4A, Table S5**). For example, several members of the CCR4-NOT and NuA4 complexes interacted with the *Phsp-4::GFP* promoter (**Figure 4A-B, Figure S20 in File S1**). Five of the CCR4-NOT complex members showed an increase in *Phsp-4::GFP* expression when knocked down, however, RNAi of one member, *ntl-4*, decreased reporter expression. Interestingly, this component is a ubiquitin-ligase that may play more of an overall regulatory function or may function outside of the CCR4-NOT complex as suggested previously(Halter et al. 2014). Additionally, each of the nine CCR4-NOT complex components interacted with at least one of the tested promoters, but no two component knockdowns had the same interaction profile (**Figure 4B**). For the NuA4 complex, six components, *ekl-4, epc-1, gfl-1, mys-1, ruvb-1*, and *trr-1*, activated *Phsp-4::GFP* upon depletion (**Figure S20 in File S1**). While NuA4 shares some components with both the SAGA and SWR1-C complexes, of the components that activate *Phsp-4::GFP* upon knockdown, only *trr-1* is shared with SAGA complex while none are shared with SWR1-C (**Table 1**).

**Figure 4.**
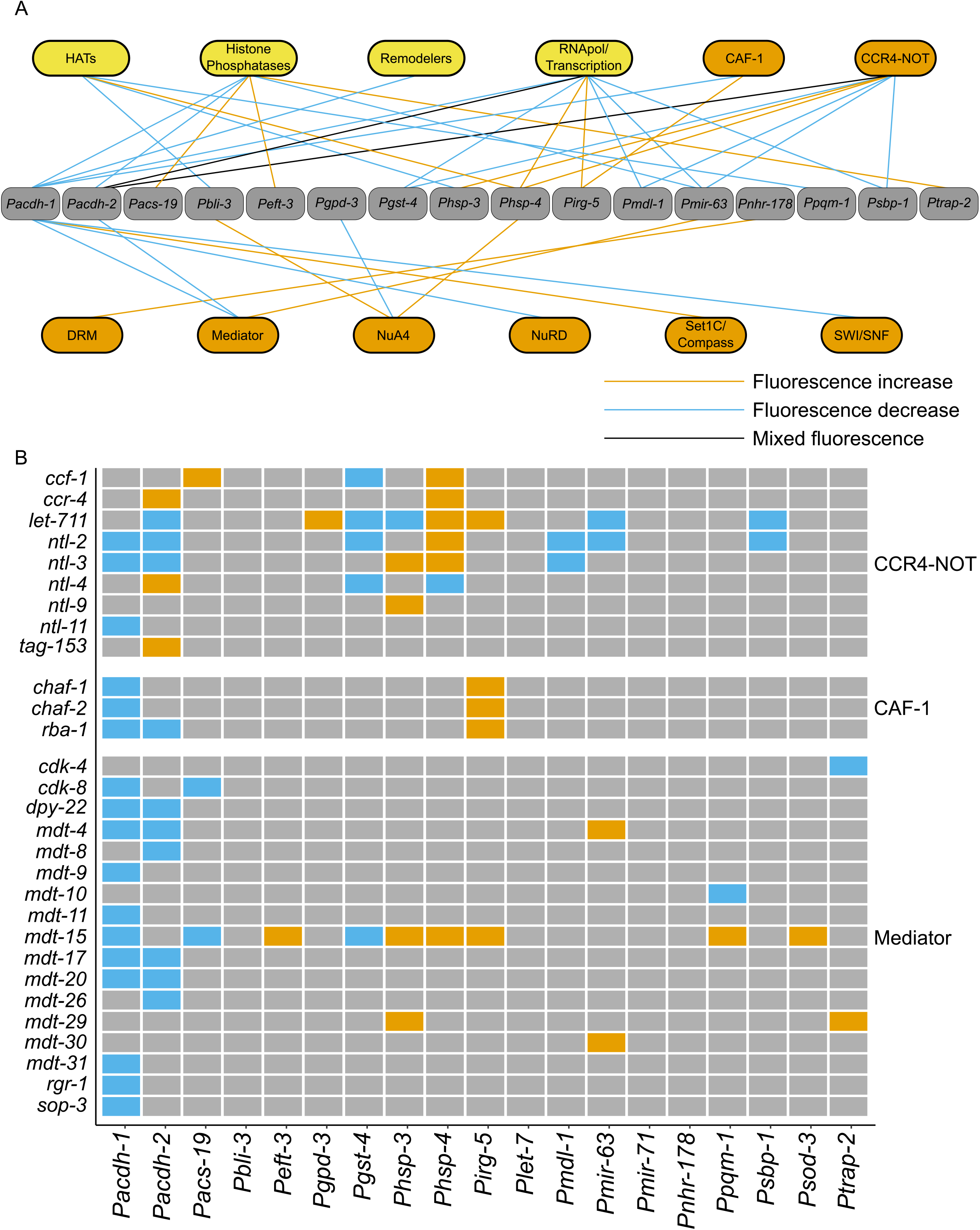
Enrichments and interaction profiles of CF complexes. (A) Hypergeometric enrichments between CF categories (yellow) and complexes (orange) that regulate the promoter reporters. Categories, complexes, and promoters without enrichments are not included. (B) Interaction profiles for CCR4-NOT, CAF-1, and Mediator complexes. Orange boxes represent increases in fluorescence (repressing interactions) and blue boxes represent decreases in fluorescence (activating interactions). For the Mediator complex, components that did not affect any of the promoters when knocked down by RNAi were not included. See also **Figure S20**.

In our dataset, CF complexes did not act uniformly. The CAF-1 complex had the most uniform effects where two members, *chaf-1* and *chaf-2*, showed the same interaction profile: their knockdown decreased fluorescence with the *Pacdh-1::GFP* reporter, and increased fluorescence with the *Pirg-5::GFP* reporter (**Figure 4B**). The third component, *rba-1*, shared those two interactions but its knockdown also decreased *Pacdh-2::GFP* expression. The nucleosome remodeling and deacetylase complex (NuRD) was also relatively uniform, with three components interacting with *Pacdh-1::GFP*, but *dcp-66* had two other interactions (**Figure S20 in File S1**). Additionally, three Mediator components, *dpy-22, mdt-17*, and *mdt-20*, have identical interaction profiles showing specificity for the *Pacdh-1* and *2* promoters. Knockdown of the well-studied Mediator component *mdt-15* affected nine promoters and exhibited both activation and repression of promoter activity when knocked down. However, only five of those promoters were affected by depletion of other Mediator components (**Figure 4B, Figure S20 in File S1**).

### Activation of *Pacdh-1* uses different TFs but common CFs

The acyl-CoA dehydrogenase ACDH-1 is the first enzyme of the propionate shunt, an alternative breakdown pathway of this short chain fatty acid that is transcriptionally activated when the canonical, vitamin B12-dependent pathway is genetically or nutritionally perturbed (Bulcha et al. 2019; MacNeil et al. 2013; Watson et al. 2013; Watson et al. 2014; Watson et al. 2016). As shown above and in previous studies, *acdh-1* promoter activity is affected by the knockdown of many CFs, TFs, and metabolic genes (**Figure 3E**)(Bhattacharya et al. 2022; MacNeil et al. 2015). We also previously found that the *acdh-1* promoter is activated in response to three specific metabolic perturbations. As mentioned above, bacterial diets low in vitamin B12 confer reduced flux through the canonical B12-dependent propionate breakdown pathway in *C. elegans* and activates *acdh-1* expression (Watson et al. 2016). Low dietary vitamin B12 also reduces flux through the Methionine/S-adenosylmethionine (Met/SAM) cycle, and mutations in Met/SAM cycle genes also increase *Pacdh-1::GFP* expression (Giese et al. 2020). Because both these activation mechanisms of *acdh-1* expression are caused by low vitamin B12; we have termed activation of *acdh-1* by excess propionate and low Met/SAM cycle activity as B12-Mechanism I and B12-Mechanism II, respectively (Giese et al. 2020). Perturbation of succinate dehydrogenase, which is also known as complex II of the electron transport chain, also activates *Pacdh-1::GFP* (Bhattacharya et al. 2022). To the best of our knowledge, low dietary vitamin B12 does not result in complex II dysfunction. Therefore, we refer to this mechanism of *acdh-1* activation as Mechanism III (**Figure 5A**). While the TFs that mediate the response to each of these have been studied in some detail, the CFs involved remain unknown.

**Figure 5.**
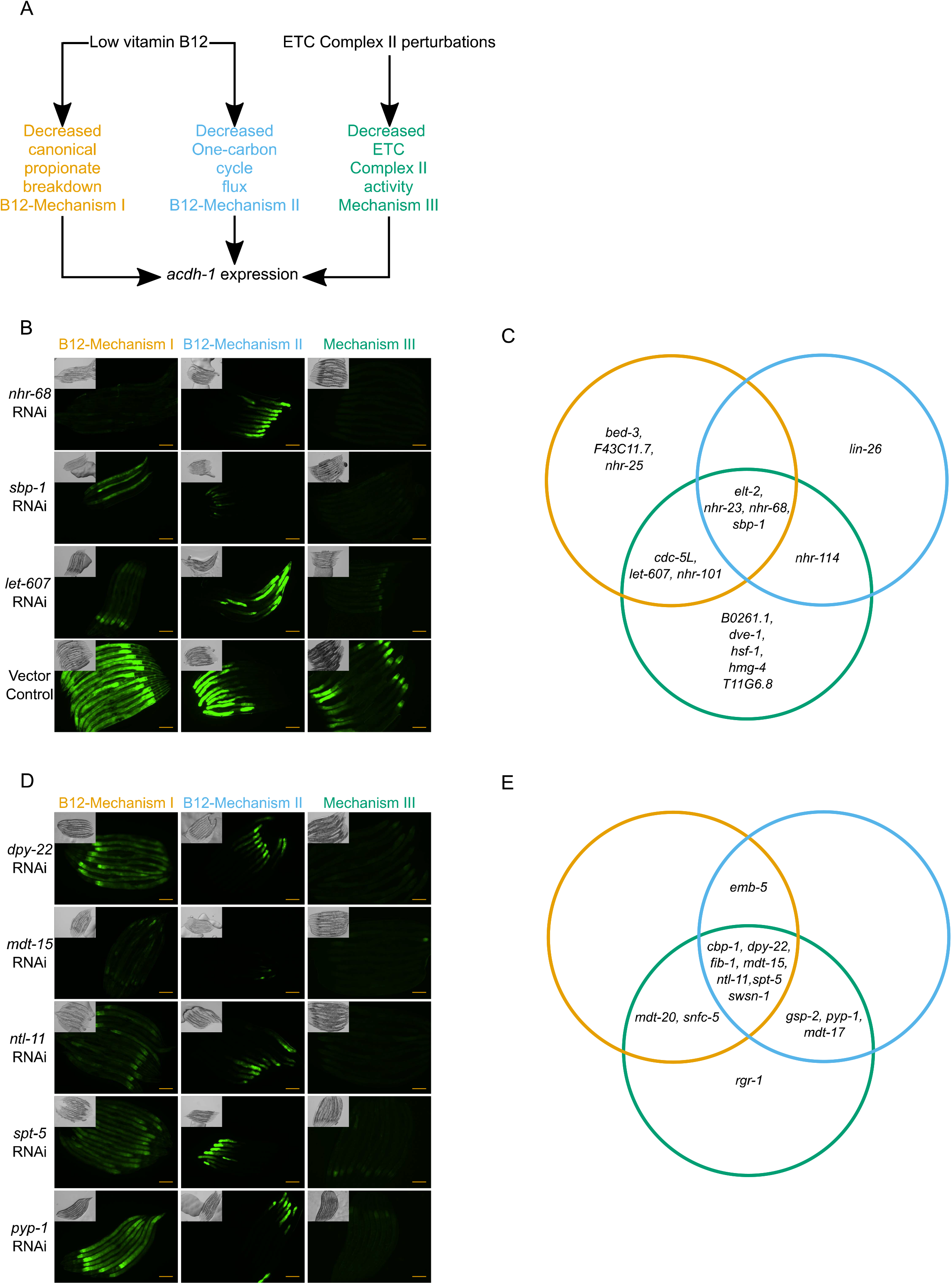
Different mechanisms that activate *Pacdh-1* use different TFs and CFs. (A) Cartoon of the three mechanisms activating *Pacdh-1::GFP*. (B) Examples of TFs that affect *Pacdh-1::GFP*. Photos of all TFs involved in at least one mechanism are provided in **Figure S21**. (C) Venn diagram of all TFs involved in the three mechanisms. Colors are as in A. (D) Examples of CFs involved in the regulation of *Pacdh-1::GFP*. Photos of all CFs that regulate *Pacdh-1::GFP* are provided in **Figure S22**. (E) Venn diagram of all CFs involved in the three mechanisms. Colors are as in A. Scale bar=100μM.

Previously, we identified forty-nine TFs that, when knocked down by RNAi, repress *Pacdh-1::GFP* under standard growth conditions (MacNeil et al. 2015). We asked which of these TFs participate in each of the three mechanisms of *Pacdh-1* activation. We found that four TFs, *elt-2, nhr-23, sbp-1* and *nhr-68*, are involved in all three activation mechanisms, while others, such as *let-607*, appear to function in only one or two of the mechanisms (**Figure 5B-C, Figure S22 in File S1**). Because analyzing B12-Mechanism II uses an *nhr-10* mutant, we did not place it in our diagram, but we note that it is required for both B12-Mechanism I and Mechanism III (**Figure S22 in File S1**).

We performed the same analysis for CFs that affected *Pacdh-1::GFP* expression, where we determined which of the CFs found to interact with this promoter in the primary screen contributed to each of the three activation mechanisms (**Figure 5D, Figure S23 in File S1**). Remarkably, in contrast to TFs, several CFs that were used for any of the activation mechanisms were involved in all three mechanisms of *Pacdh-1* activation (**Figure 5E**). Of the fourteen CFs used by at least one mechanism, seven were used in all three. Interestingly, there were five components of the Mediator complex required for Mechanism III activity: *dpy-22/mdt-12, mdt-15, mdt-17, mdt-20*, and *rgr-1/mdt-14*. Several TFs that regulate *Pacdh-1*, including SBP-1 and NHR-10, physically interact with MDT-15(Arda et al. 2010). These results indicate that different combinations of TFs and CFs function together to induce *acdh-1* expression in response to different metabolic perturbations. Further, the observation that not all TF and CF knockdowns that activate *acdh-1* under standard conditions were found to act in any of the three mechanisms indicates that there could be additional mechanisms of *acdh-1* induction.

### Three CBP/p300 paralogs with only partially overlapping domains all regulate promoter activity

As discussed above, the CBP/p300 ortholog *cbp-1* regulates the greatest number of the tested promoters of the CFs tested. This CF has been extensively studied in many eukaryotic model systems and was found here to function as both a transcriptional activator and repressor (Boija et al. 2017; Hunt et al. 2022). In early *C. elegans* development, *cbp-1* is required for proper cell fate decisions (Shi and Mello 1998). The *C. elegans* genome also encodes two shorter CBP/p300 homologs, *cbp-2* and *cbp-3*. All three *C. elegans cbp* genes are located within a 160 kb stretch of chromosome III and are likely the result of partial gene duplications, as they share sequence homology (**Figure 6A**). Both *cbp-2* and *cbp-3* are annotated as pseudogenes, although both are expressed at the mRNA level in the intestine (Cao et al. 2017; Mitrovich and Anderson 2005; Shi and Mello 1998). We examined InterPro annotations for protein domains and found that CBP-1 has four zinc finger domains, a RING domain, a KIX domain, a bromodomain, and a large HAT domain (**Figure 6B**). However, CBP-2 only contains the first zinc finger domain and the KIX domain, and CBP-3 only contains the first zinc finger domain (**Figure 6B**). By these annotations, neither CBP-2 nor CBP-3 would have acetyltransferase activity on their own. In our screen, all three *cbp* paralogs interacted with a subset of promoters. However, we noticed that the *cbp-2* clone we obtained from the ORFeome library (Rual et al. 2004) actually contained the *cbp-3* sequence. The sequences of all three *cbp* genes closely align, and the 5’ and 3’ ends of *cbp-2* and *cbp-3* are particularly similar (**Figure 6A**). We created a new *cbp-2* RNAi clone that targets a more unique sequence within the *cbp-2* mRNA and verified that both this and the RNAi clone for *cbp-3* do not affect *cbp-1* expression (**Figure 6C**). We then tested the new *cbp-2* RNAi clone together with the clones for *cbp-1* and *cbp-3* to determine whether all three paralogs interact with the same 14 promoters that were affected by *cbp-1* RNAi (**Figure 6D**). Although neither *cbp-2* nor *cbp-3* affected all fourteen reporters, both did regulate multiple promoters. While these results do not unveil the extent to which the *cbp* paralogs act in *C. elegans* genetic regulation, these findings indicate that *cbp-2* and *cbp-3* may produce functional proteins and may not be pseudogenes.

**Figure 6.**
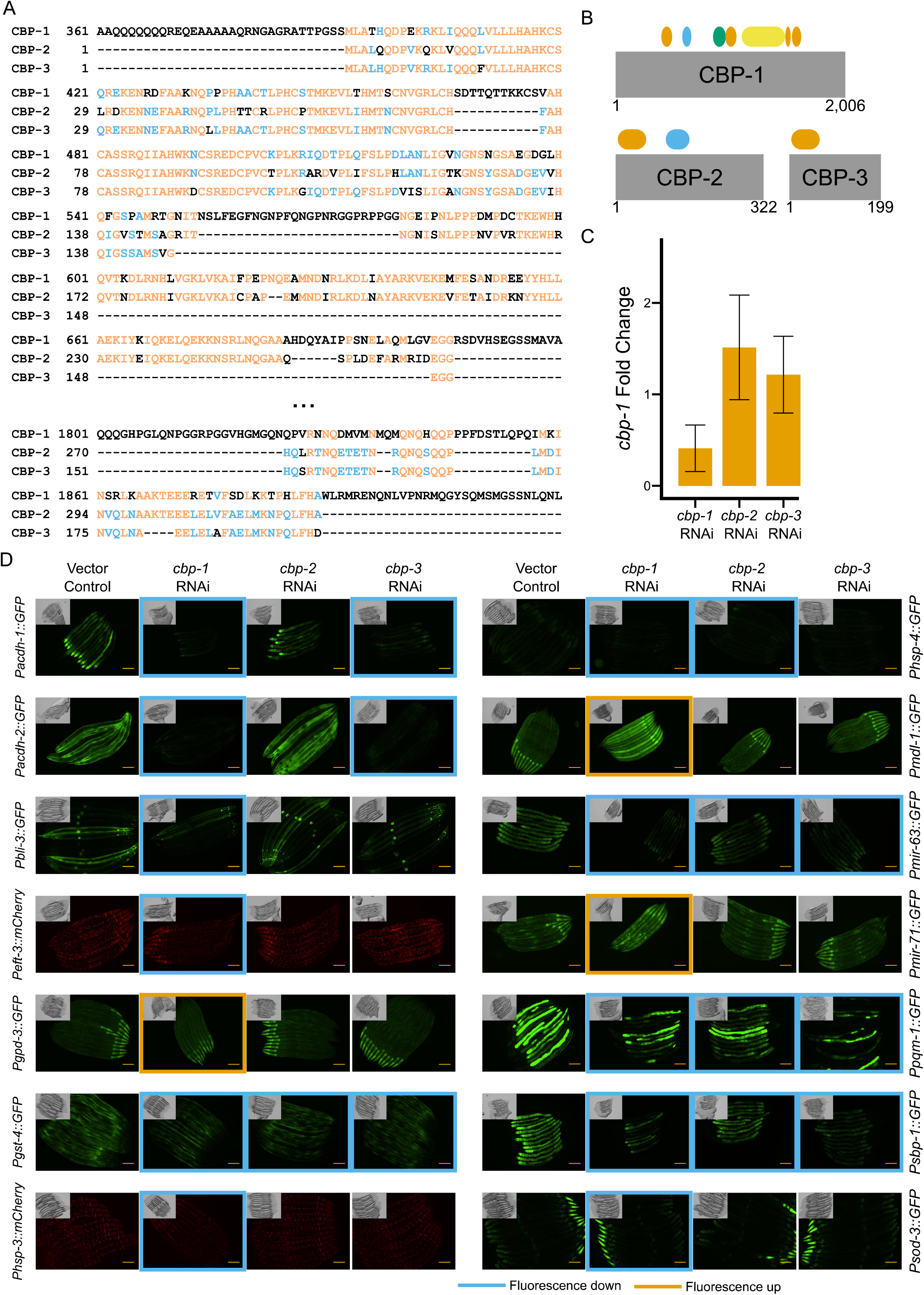
*cbp-1, cbp-2*, and *cbp-3* are functional genes. (A) Sequence alignment of CBP-1, CBP-2, and CBP-3. Black letters are unique to one protein, blue letters are common to two proteins, and orange letters are common to all three proteins. Portions of *cbp-1* sequence that do not align to *cbp-2* or *cbp-3* are not shown. (B) Protein domains encoded in CBP-1, CBP-2, and CBP-3. Orange represents zinc finger domains, blue represents KIX domains, green represents a bromodomain, and yellow represents a HAT domain. Numbers below indicate amino acid number. (C) Fold change of *cbp-1* upon *cbp-1, cbp-2*, and *cbp-3* knockdowns compared to vector RNAi. Bars represent the mean of three biological replicates, error bars represent one standard deviation. (D) Photos of fourteen strains with vector control, *cbp-1, cbp-2*, and *cbp-3* RNAi. Strains that do not interact with *cbp-1* are not shown. Scale bar=100μM.

## DISCUSSION

In this study we generated a comprehensive RNAi resource for *C. elegans* CFs and examined how the depletion of each of these affects promoter activity in the *C. elegans* intestine. There are clear advantages to our promoter-centered approach, including the ability to comprehensively test the effects of regulators such as CFs, the ability to specifically examine changes in one tissue in living animals, and the knowledge of the DNA element through which the effect on reporter expression occurs, in this case usually ∼2kb gene promoters. However, there are several distinct technical and conceptual disadvantages as well. Technical challenges include the fact that high-throughput RNAi screens can be noisy, *i*.*e*., they miss interactions. This is because the screens are done visually, because not all promoter reporters are integrated into the genome, resulting in mosaicism of fluorescence, and because some promoters drive low levels of fluorescent protein expression, which makes it more difficult to observe reductions in fluorescence upon RNAi of a regulator. A notable conceptual disadvantage includes the focus on a single tissue only during development.

Overall, we observed very few changes with the knockdown of methyltransferases, demethylases, HDACs, remodelers, or TBP-associated proteins. Instead, the majority of expression changes occurred through depletion of HATs, RNA pol-II-associated factors or Mediator components. However, because we only used 19 promoters, it remains to be determined how generalizable these observations are. Several studies have revealed that many CFs regulate specific genes in other organisms(Haberle et al. 2019; Lenstra et al. 2011; Neumayr et al. 2022). Since we also find that many CFs act specifically on some promoters without detectable effects on others, these findings together indicate that many CFs act in a highly gene-specific manner.

The CF that interacted with the greatest number of promoters was *cbp-1*, which affected 14 of the 19 promoters. While its ortholog p300/CBP has been mostly known for its role in transcriptional activation through its acetyltransferase activity, it has recently been shown to have repressive functions independent of this enzymatic activity as well(Hunt et al. 2022). Two paralogs of *cbp-1, cbp-2* and *cbp-3*, were previously annotated as pseudogenes. However, we found that independent depletion of either of these genes affected promoter activity. Neither of the proteins encoded by these homologs are predicted to have a HAT domain, indicating that they act by other mechanisms. Future work will be needed to further characterize the biological function of these two genes and how they affect gene expression.

We previously used the same 19 reporter strains to identify TFs and metabolic genes that affect promoter activity(Bhattacharya et al. 2022; MacNeil et al. 2015). While depletion of CFs or TFs frequently decreased promoter activity, knockdown of metabolic genes generally resulted in increased promoter activity(Bhattacharya et al. 2022; MacNeil et al. 2015)(this study). In the TF study, we found that many TFs affected promoter activity indirectly, *i*.*e*., without apparent physical binding. Since metabolism and gene expression frequently influence each other(Carthew 2021; Giese et al. 2019; Li et al. 2018; Watson et al. 2015), we hypothesized that metabolism may connect TFs that indirectly regulate promoter activity to TFs that both bind and regulate promoter activity. Similarly, we reasoned that because CFs often connect metabolism and gene regulation, for instance by modifying DNA or histones with metabolites such as methyl and acetyl groups(van der Knaap and Verrijzer 2016), we may be able to place these transcriptional regulators in the context of larger gene regulatory networks comprising promoters, TFs, metabolic genes, and CFs. Previously, we used epistasis-based nested effects modeling to organize the TF-based gene regulatory network into a hierarchy that depicts the regulatory ‘flow of information’(MacNeil et al. 2015; Markowetz et al. 2007). Here, we were unable to use the same approach to connect TFs with CFs and metabolic genes. This could be because of the size of the datasets that need to be combined, because the data are relatively noisy, and/or because the output of nested effects modeling is difficult to navigate. We propose that it may be more feasible to build regulatory networks for individual promoters, one-at-a-time, to incorporate different types of regulators.

We explored the concept of combining different types of regulators for individual promoters with the *acdh-1* promoter, which is affected by the greatest number of TFs, CFs and metabolic genes and which we have previously studied in more detail(Bhattacharya et al. 2022; Bulcha et al. 2019; Giese et al. 2020; Watson et al. 2013; Watson et al. 2014)(this study). There are currently at least three known mechanisms of *acdh-1* activation, some of which we have studied in detail, and some of which remain to be elucidated further. First, *acdh-1*, like all 19 promoters, is activated by the intestinal master regulator *elt-2*. This GATA TF activates genes, including those encoding TFs, and resides at the top of the regulatory hierarchy (MacNeil et al. 2015). The SREBP ortholog, *sbp-1*, induces additional TFs and also activates *acdh-1*. These TFs include *nhr-68*, which acts in a type I coherent feed-forward loop with *nhr-10* to activate *acdh-1* in response to sustained propionate accumulation (Bulcha et al. 2019; Ding et al. 2015). SBP-1 also activates *nhr-114*, which activates *acdh-1* in response to low Met/SAM cycle flux(Giese et al. 2020). Here, we found that different CFs do not appear to provide additional specificity to *acdh-1* activation in response to different metabolic perturbations. Therefore, the molecular mechanisms by which TFs and CFs converge on the *acdh-1* promoter under different metabolic perturbations, and how these regulatory effects are insulated from each other remain to be elucidated, for instance by using chromatin-immunoprecipitation-based methods. Finally, using more genome-scale methods such as perturb-seq may shed further light on the interplay between TFs, CFs, and metabolic genes.

## FIGURE LEGENDS

**File S1**. Supplemental Figures S1-S22.

**Table S1**. CFs encoded in the *C. elegans* genome.

**Table S2**. Hypergeometric enrichments for essentiality among CF classes, CF binding domain-containing proteins, and for CFs, TFs, and metabolic genes.

**Table S3**. Description of *C. elegans* strains used in the primary screen.

**Table S4**. Interactions between CF RNAi strains and the 19 promoter reporters.

**Table S5**. Hypergeometric enrichments for interactions between CF classes or complexes and promoter reporters.

**Table S6**. Oligonucleotides used in this study.

## AUTHOR CONTRIBUTIONS

BBH and AJMW conceived the study and annotated *C. elegans* cofactors. BBH performed most experiments and analyses with help from SN (intestinal gene expression).

## DATA AVAILABILITY STATEMENT

All interaction data generated in this study are available in the article and associated supplementary files. The CF RNAi library is available from the corresponding author upon request.

## ACKNOWLEDGEMENTS

We thank the present and previous members of the Walhout lab, particularly Sushila Bhattacharya, Amy Holdorf, Yomari Rivera, Akshaye Shah, and Caryn Navarro, for technical support, discussion, and comments on experiments and the manuscript.

